# Gaze-related functions driving gaze anchoring in reaching

**DOI:** 10.1101/2025.08.20.671334

**Authors:** Jolande Fooken, Nethmi H. Illamperuma, Jason P. Gallivan, J. Randall Flanagan

## Abstract

When reaching to a target, we typically direct our gaze to the target throughout the reach. If a secondary saccade target is presented during the reach, and we are instructed to shift our gaze to this target, the gaze shift is generally delayed until the reach target has been attained—a phenomenon known as gaze anchoring. Here, we compared gaze anchoring in human participants when reaching to a visual target versus a visual-haptic target providing force feedback upon contact. We also examined gaze anchoring in a bimanual context in which participants reached with their right hand and were instructed to both shift their gaze and move their left hand to the secondary target as soon as it appeared. We found that, during the reaching movement, gaze was anchored to the target for both visual and visual-haptic targets. In the visual-haptic condition, gaze appeared to be anchored during the initial directing phase of the reach during which peripheral vision directs the hand to the vicinity of the target. In the visual condition, gaze appeared to be anchored during both the directing phase and subsequent guiding phase during which central visual guides the hand. In two-handed reaching, gaze anchoring was observed but anchoring did not extend to the left hand, which started moving before the eyes. Overall, our findings indicate that the timing of eye and hand movements in object manipulation is linked to the function of target fixations.

**New & Noteworthy:** When reaching to a visual target, gaze is typically anchored to the target even when a competing saccade target appears. We show that anchoring also occurs for visual–haptic targets but is limited to the initial phase of the reach, likely because vision is not required to confirm target attainment. In contrast to gaze, anchoring is not observed for the non-reaching hand, which can begin moving to a competing target as soon as it appears.

## Introduction

Many natural action tasks, such as cooking, require a sequence of coordinated eye and hand movements. In goal-directed action tasks, in which people reach for and manipulate objects, gaze commonly fixates the next object that is to be manipulated at the start of the reach, and shifts to the next target of interest around the time the hand arrives at the current target (Ballard et al. 1992; Bowman et al. 2009; Epelboim et al., 1995; Wilmut et al. 2006). For example, in block stacking, gaze shifts from a given block to its placement location around the time the hand contacts the block (Flanagan and Johansson 2003).

Fixating the target serves several functions, including ‘directing’ and ‘guiding’ the hand to the target and ‘checking’ goal completion (Land et al. 1999; Land 2006; Land and Hayhoe 2001). In the context of reaching, directing refers to the use of peripheral vision and gaze-related signals—including proprioceptive signals of the eye and/or efference copy of motor commands—to control the reaching movement towards a foveated target (Bridgeman and Stark 1991; Goettker et al. 2020; Goodale et al. 1986; Prablanc et al. 1986). During the directing phase of the reach, peripheral vision of the hand can be used to rapidly (∼150 ms) and automatically correct for reach errors (Brenner and Smeets 1997; de Brouwer et al. 2018; Gallivan et al., 2016; Paillard 1996; Sarlegna et al. 2003; Saunders and Knill 2003, 2004). Guiding refers to the use of central (i.e., parafoveal) vision to control the hand movement as it approaches and contacts the reach target (Johansson et al. 2001). During the guiding phase of the reach, central vision of the hand and target can be used to adjust the hand via relatively slow feedback loops. Finally, checking refers to the use of central vision to confirm that contact between the hand and target has been achieved (Säfström et al. 2014). Many studies of goal-directed reaching have used purely visual targets, in which case central vision is required to guide the hand and check goal completion. However, when reaching to physical objects, haptic information can often be used to guide the hand and check goal completion.

In real-world manipulation tasks, there is often competition for gaze resources between targets of action and events in the environment, such as when a brightly coloured bird appears in the peripheral vision of a birdwatcher who is reaching towards their foveated binoculars. In a series of studies, Neggers and Bekkering (2000, 2001, 2002) examined such competition using a task in which a secondary saccade target was visually cued while participants reached towards a foveated reach target. The visual target was cued at different times during the movement and the participant was instructed to shift their gaze to the saccade target as soon as it was cued. When the cue was presented during the reach, gaze remained ‘anchored’ at the reach target until around the time the hand arrived. As a consequence, the latency of the saccade—or saccadic reaction time—increased with the remaining duration of the reach movement after the cue. In the current study, we aimed to link the timing of saccades and reaching movements to the functional demands of gaze.

Previous work on gaze anchoring during target-directed reaching has focused on tasks in which the participant moves their hand to visual targets (Abekawa et al. 2021; Neggers and Bekkering 2000, 2001). In this scenario, we would expect gaze to support both directing and guiding the hand to the target in addition to checking goal completion. However, it is unclear whether gaze anchoring is taking place in the directing phase of the reach, or whether anchoring is only taking place towards the end of the reach movement (i.e., during guiding and checking). Importantly, peripheral vision can be effectively used to direct the hand when fixating a gaze target that is displaced from the reach target (de Brouwer et al. 2017, 2018; Neggers and Bekkering 2002). Thus, it is plausible that, when a secondary saccade target is presented during the reach, participants maintain gaze at the reach target because it is critical for the forthcoming guiding and checking phases, rather than for directing per se.

To examine whether gaze anchoring occurs during directing, we examined eye-hand coordination during a reaching task (Fig. 1A) in which the participant moves a hand-held handle, represented as a cursor, with their right hand to a target. In the visual-haptic condition, contact forces between the cursor and the target are simulated by applying forces to the hand via the handle. In the standard visual condition, no contact forces are simulated.

**Figure 1.**
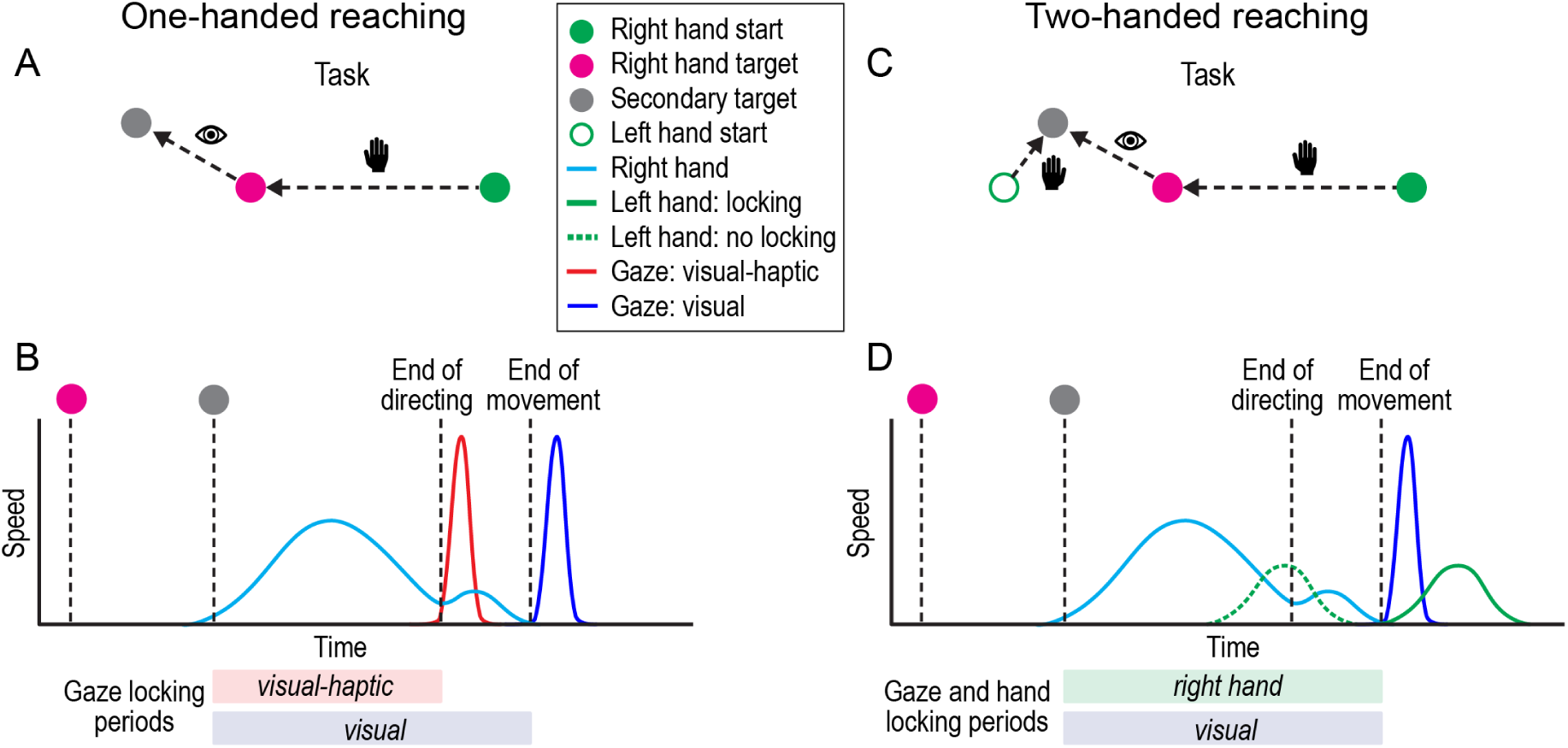
Tasks and predicted results. (A) In the one-handed reaching task, the participant is instructed to shift their gaze from the reach target to a secondary saccade target, which is cued at different times during the right-hand reach. In the visual-haptic condition, both visual and haptic feedback provides confirmation of reach target attainment whereas only visual feedback does so in the visual condition. (B) Schematic showing hand and gaze speed over time with predicted periods of gaze-locking in the visual-haptic and visual conditions when the secondary gaze target is cued at the beginning of the movement. (C) In the two-handed task, the participant is instructed to move their gaze and left hand to the secondary (saccade + left hand) target, which is cued at different times during the right-hand reach to the initially foveated reach target. (D) Schematic showing right and left hand and gaze speed over time with predicted periods of gaze- and hand-locking when the secondary gaze target is cued at the beginning of the movement.

We expected that in the visual-haptic condition, gaze would not be required for guiding because haptic feedback can be used to guide the hand to the target and confirm target attainment. Thus, if gaze anchoring occurs during directing, we should find that gaze is locked to the target as the hand moves towards the target but is released (i.e., is able to shift to the secondary target) at the end of the directing phase—and the start of the guiding phase—when the hand arrives close to the target. Figure 1B illustrates the predicted periods of gaze locking in the visual and visual-haptic conditions when the secondary gaze target is presented at the start of the movement.

In addition to testing whether gaze anchoring occurs during the directing phase of visually guided reaches, we asked whether the timing of gaze anchoring would be affected if the saccade target also became a hand movement target. In a two-handed version of our reaching task (Fig. 1C), participants used their right hand to reach to a foveated visual reach target, just as in the one-handed task, and were instructed to move both their gaze and their left hand to a secondary target as soon as it was cued. Note that we only included a visual condition in the two-handed task. Figure 1D illustrates the predicted periods of gaze and left-hand locking when the secondary gaze + left hand target is presented at the start of the movement. We expected saccade latencies in the two- and one-handed tasks to be similar because the visual and gaze-related functional demands of the right-handed reach should be the same. If the left hand exhibits “hand locking” we would expect the left-hand movement to be initiated at the end of the movement (solid green trace in Fig. 1D). However, if hand locking is not observed, we would expect that the onset of the left-hand movement to be linked to the presentation of the secondary target and be delayed by a standard reaction time period (dotted green trace in Fig. 1D).

## Methods

### Participants

A total of twenty-eight individuals participated in the study (18-51 years of age; mean age 25, 4 left-handed). Participants were primarily recruited from the Queen’s University undergraduate and graduate student population. All participants were eligible for monetary compensation ($10 per hour) or course credits towards a psychology course. Each participant provided written consent prior to participation and received a debriefing once the experiment was completed. Participants were required to be 18 years of age or older, and have no history of psychological, neurological, or eye disease. The experiment was approved by the Queen’s General Research Ethics Board (TRAQ #: 6003707) and complied with the Declaration of Helsinki.

### Apparatus

Participants were seated at a desk with their head placed on a mounted chin and forehead rest in front of a vertical monitor (70 × 39.5 cm in size; 1920 × 1080 resolution) on which the visual stimuli were displayed. Eye movements were recorded using a desktop mounted eye tracker (EyeLink 1000; SR Research, Ltd., Kanata, ON, Canada). Unimanual and bimanual reaching movements were performed in a horizontal plane using the handles of a robotic manipulandum (End-Point robot, KINARM, Kingston, ON, Canada). The position of the hand was represented as a cursor on the vertical monitor with forward and rightward hand movements mapped onto upward and rightward cursor movement (as with a standard mouse). There was a 1:1 relationship between hand movement displacement and cursor movement displacement, i.e., a 10 cm movement of the hand resulted in a 10 cm movement of the cursor. The viewing distance from the right eye to the monitor was 37 cm, such that a 1 cm displacement on the monitor corresponded to 1.5 visual degrees.

### Stimuli and Procedure

Three conditions (one-handed visual, one-handed visual-haptic, and two-handed visual) were examined with 108 trials per condition. Fourteen participants performed the one- and two-handed visual reaching conditions, and 14 separate participants performed the visual-haptic reaching condition. Note that this sample size is similar to those we have used in our previous studies of eye-hand coordination (e.g., Johansson and Flanagan, 2001; Flanagan and Johansson, 2003; Fooken et al., 2024b). Before each block, participants received training to become familiar with the equipment and to ensure consistent right-handed reaching speed within and across participants. Specifically, participants performed 24 right-hand reaches, in which no secondary saccade target was shown. After each reach in the training, participants received visual feedback in form of a horizontal line that would be above (too slow) or below (too fast) the target line (movement duration of 500 ms). Following training, the eye tracker was calibrated, and participants were asked to keep their head in the chin and forehead rest throughout each block.

The KINARM correct for buffering delays in the display system by using the current velocity of the hand (e.g., robot handle) to predict—and display—the position of the hand ∼50 ms into the future. The delay between the time an event code was sent and the time visual stimuli appeared was on average 53 ms (range: 41-64 ms).

#### One-handed reaching experiment

In this experiment, two groups of participants perform the one-handed reaching task. Participants in the visual condition did not experience any forces when reaching. In contrast, participants in the visual-haptic condition experienced a force on the hand (via the handle of the robotic manipulandum) when the cursor, controlled by the hand, contacted the reach target. Specifically, as soon as the outer part of the cursor overlapped with the outer part of the reach target, the robot generated an elastic force (with a stiffness of 15 N/cm) that ‘pulled’ the handle to the centre of the reach target and held it there until the end of the trial. Whereas participants could feel the force upon contact, they did not explicitly become aware of the assistive pull.

In both conditions, participants used their right hand to move the robot handle (Fig. 2A). To initiate a trial, participants had to move the blue-coloured cursor (0.5 cm in diameter) inside the start position. Then, three grey hollow circles (0.8 cm in diameter), serving as saccade targets, and one pink-coloured hollow circle (0.8 cm in diameter), serving as the reach target, appeared. Participants were required to keep their gaze on the pink-coloured hand target and their hand at the start position for a randomly jittered time period of 1-1.5 seconds. After the fixation period, the pink-coloured hand target filled in, prompting participants to initiate a reaching movement from the start position to the hand target. The distance from the start position to the hand target was 15 cm. During this reaching movement, one of the three grey saccade targets filled in, prompting participants to move their gaze from the hand target to the cued saccade target as soon as possible. The horizontal saccade target was 10 cm away from the hand target, and upper and lower targets had a horizontal offset of 7 cm and a vertical offset of ± 4 cm (i.e., ± 52° from the horizontal).

**Figure 2.**
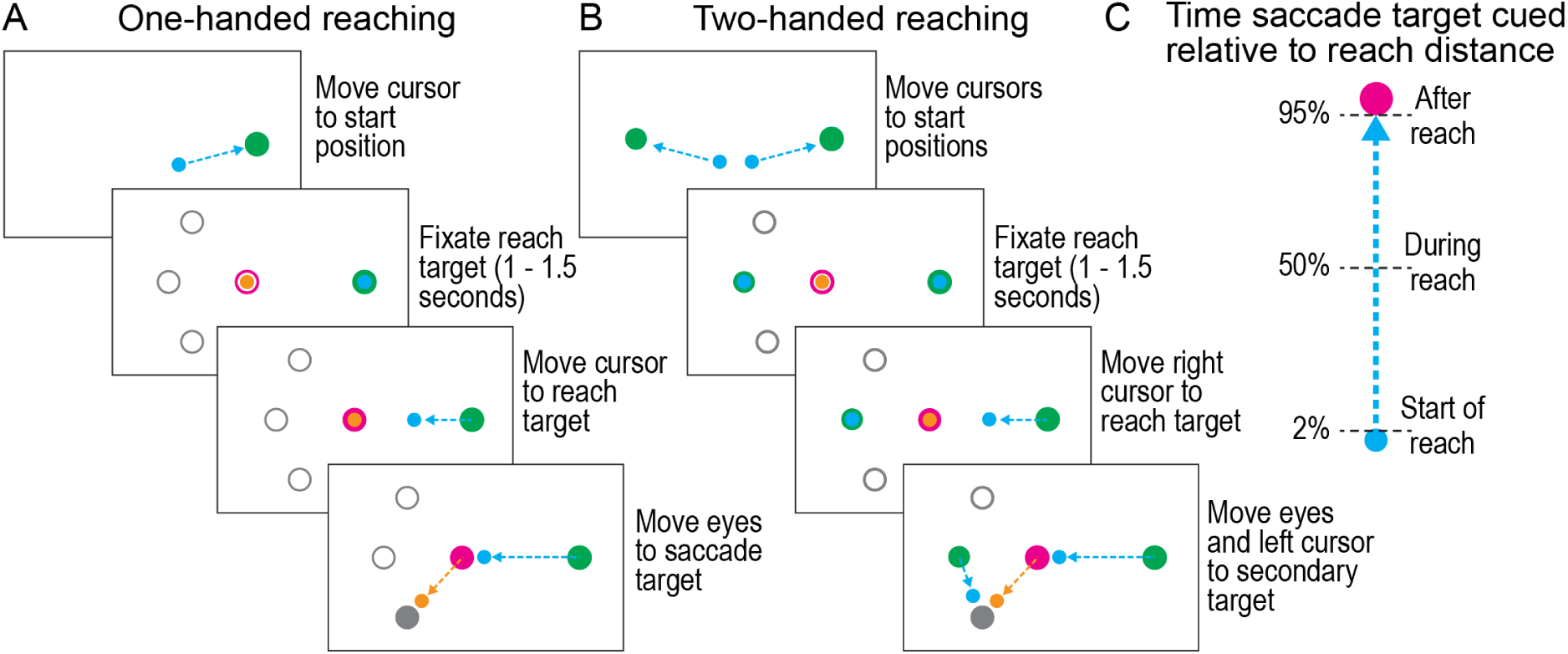
Experimental conditions and task procedure. (A) In the one-handed reaching task, participants moved the cursor corresponding to their right-hand position to the start location. Participants were instructed to fixate a hollow pink reach target and, once it filled in, to reach from the start to the reach target using their right hand. While reaching, one of three hollow grey saccade targets filled in, cueing the participant to move their eyes to the saccade target as soon as possible. (B) In the two-handed reaching task, participants moved both cursors, indicating their left and right hand, to the respective start locations. Participants were instructed to fixate a hollow pink reach target and keep their left hand at the left start location. Once the pink reach target filled in, participants moved their right hand to the reach target. While reaching, one of two hollow grey saccade targets filled in, cueing the participant to move their eyes and their left hand to the combined saccade and left-hand target as soon as possible. One group of 14 participants completed the one- and two-handed reaching tasks with visual targets, where ‘contact’ between cursor and reach target was visual only. A second group of 14 participants performed the one-handed task with visual-haptic targets with simulated contact forces between the cursor and reach target. (C) Saccade targets or combined saccade and left-hand targets were cued at the start, during, or after the right-handed reaching movement.

The saccade target was cued in every trial and could appear at one of three possible times during the reaching movement—when the hand had moved 5% (start of reach), 50% (during reach), or 98% (end of reach) of the distance from the start to the hand target (Fig. 1D)—inciting one of the grey circles to fill in. The position and time at which the eye target was cued was randomized between trials. The trial finished once the hand arrived at the hand target, and the cursor had to be within the hand target for 250 milliseconds for the trial to successfully finish. During the experiment, participants were notified if they moved their hand before the reaching movement was cued (pink target filled in) or moved their gaze before the saccade target was cued (one of three grey targets filled in). Trials in which participants moved their eyes or hands too early were immediately repeated.

#### Two-handed reaching experiment

In the two-handed visual condition, participants used both hands to move the robot handles (Fig. 2B). The hand start positions, hand target, and saccade targets were the same size as in one-handed reaching. The right start position was 15 cm to the right of the reach target, and the left start position was 9 cm leftward of the right-hand reach target. At the start of each trial, participants moved the left and right robot handle to place the blue-coloured cursor at the left and right start position, respectively. Two grey hollow circles (the upper and lower saccade targets from the one-handed reaching condition) and one hollow, pink coloured right-hand target appeared. Participants were required to keep their gaze on the right-hand reach target and their two hands at the respective start positions for 1-1.5 seconds. After the fixation period, the right-hand pink target filled in, prompting participants to initiate a reaching movement from the right start position to the right-hand target. Next, one of the two combined saccade and left-hand targets was cued at the same cueing times as in the one-handed reach condition (start of reach, halfway through reach, end of reach). Once the combined saccade and left-hand target filled in, participants were instructed to move both their left hand and their eyes to the cued eye-hand target as soon as possible. Again, participants received immediate feedback, and trials were repeated if they moved their eyes or either of their hands before the respective cue.

### Analysis

To analyze eye and hand movement data, we created custom-made routines using MATLAB (version 2023b). For hand movement analyses, we analyzed the centred x and y positions of the robotic handles that were sampled at 1000 Hz. Position data were filtered using a third order 20 Hz Butterworth filter. For all reaching movements, we determined the start of the reach, the end of the directing phase—which coincided with the start of the guiding phase—and the end of the guiding phase (i.e., the end of the movement). The start and end of the reach were defined at the times at which the hand velocity first exceeded and subsequently dropped below 5% of the peak velocity of the current reach, respectively. To estimate the time point at which the directing phase ended, and the guiding phase started, we first selected all samples during which the hand decelerated. We then differentiated this hand deceleration and found the time at which the jerk peaked. In bimanual reaching trials, we further analyzed left-hand latency. Hand latency was defined as the difference between the time at which the combined left-hand and saccade target was cued and the start of the left-hand movement, defined as the time at which the left-hand velocity first exceeded 5% of the peak velocity of the current reach.

We recorded the x- and y-position of the right eye with a sampling rate of 500 Hz. Eye position signals were filtered using a second-order 15 Hz Butterworth low pass filter. Eye velocity was determined by differentiating the eye position signal. Eye velocity samples were labelled as saccades when five consecutive samples exceeded a fixed velocity criterion of 50 cm/s. To determine saccade onsets and offsets, we found the nearest reversal in the sign of the eye acceleration signal before eye velocity exceeded the fixed velocity threshold (saccade onset), and the nearest reversal in the sign of eye acceleration after eye velocity was back below the fixed velocity threshold (saccade offset). We then calculated saccade latency relative to the time of saccade cue, and saccade latency relative to the time at which the right hand first contacted the reach target.

For the one- and two-handed reaching, one participant was excluded because they were unable to follow task instructions and moved their eyes away from the reach target before the reaching movement was cued in a majority of trials. One other participant was not included in the visual contact group because of an erroneous KINARM calibration. For the remaining 26 participants (13 in experiment 1 and 2, respectively) we excluded 717 out of 12636 trials (5.7%). Specifically, we excluded trials in which participants moved their eyes or left hand anticipatory to the cue (before the time of cue or within 100 ms of the cue, respectively; 1.6%), the eyes did not land on the cued target (1.5%), or fixation on the reach target was not maintained (2.6 %). Fixations on the reach target were not maintained due to blinks, saccades to the hand starting position(s), or to the visual scene.

We used mixed-factor and repeated measures ANOVAs to test for effects of cue time and condition. We also used linear regression to examine individual differences. A *p* value ≤ 0.05 was considered to be statistically significant.

## Results

### Gaze anchoring in one-handed reaching depends on sensorimotor feedback

In the one-handed reaching task, we aimed to link the timing of eye and hand movements to the functional demands of gaze. Participants performed right-handed reaches to move a cursor, controlled by the hand, to a stationary reach target. After, during, or at the start of the reach, a secondary saccade target filled in, cueing participants to shift their gaze from the fixated reach target to the saccade target as soon as possible. In the visual condition, participants only received visual feedback about contact (i.e., when the cursor “contacted” the target), whereas in the visual-haptic condition, participants also experienced a force upon contact.

Figures 3A and B show, for representative trials from the visual and visual-haptic conditions, eye and hand (i.e., cursor) paths and target positions in a screen-centred reference frame. Figures 3C and D show gaze and hand movement velocities as a function of time for the same trials. The time of the cue and the end times of the directing phase and movement are shown by vertical lines. In both of these trials, the top potential saccade target was cued as the saccade target just after movement started. Gaze shifted to the saccade target just after the end of the movement in the visual condition, and just after the end of the directing phase (and before the end of the movement) in the visual-haptic condition. Figures 3E and F show each participant’s mean hand movement velocity (thin lines) and the group average (thick line) as a function of time. Velocity profiles were normalized to the median durations of the directing and guiding phase in the visual condition and haptic-visual condition, respectively. The duration of the hand movement was slightly shorter (*t*(24) = 2.07; *p* = 0.05) in the visual-haptic condition (mean participants’ median reach duration: M = 603 ms, standard error: SE = 36 ms) compared to the visual condition (M = 708 ms, SE 36 ms).

**Figure 3.**
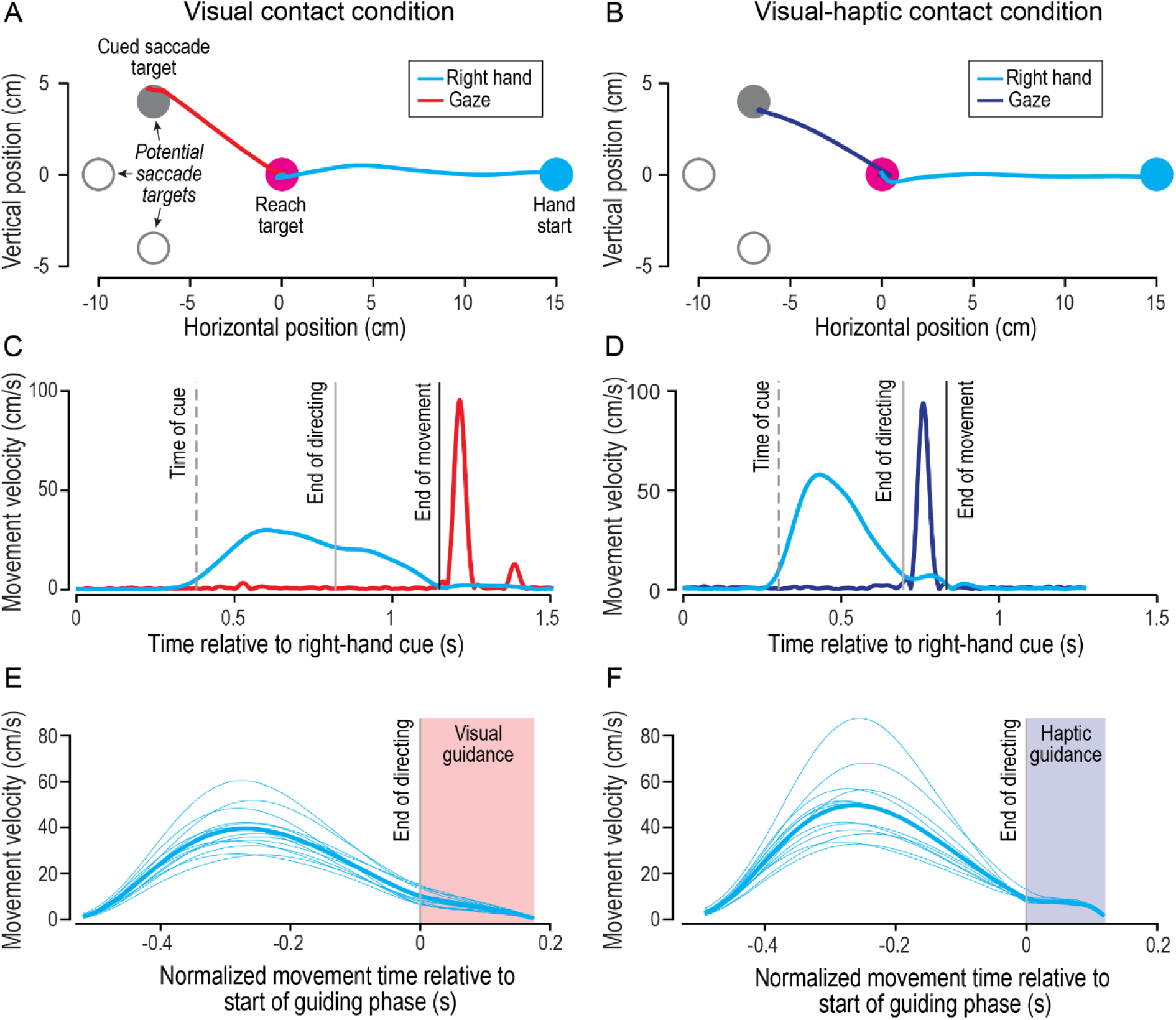
Results from the one-handed reaching task. (A, B) Gaze and hand (i.e., cursor) paths and target positions, in screen coordinates, from representative trials from the visual and visual-haptic conditions. Participants used their right hand to reach from the start position to the reach target and were instructed to shift their gaze from the reach target to cued saccade target, randomly selected from three potential saccade targets. (C, D) Gaze and hand tangential velocity profiles from the trials in A and B. The times of the cue and ends of the directing phase and movement are indicated by vertical lines. (E, F) Mean hand velocity profiles of all participants (thin lines) and averaged across the two conditions (thick lines). Velocity profiles were normalized to the median duration of the directing and guiding phase, respectively.

Depending on the position of the saccade target, the saccade from the hand target to the saccade target required either a horizontal or an oblique eye movement. We found no difference in saccade latency between horizontal and oblique targets (*t*(12) = 1.11; *p* = 0.29) and thus averaged across saccade target locations.

We used a mixed-factor ANOVA to examine the effects of the between-participants factor condition (visual vs visual-haptic) and the within-participants factor saccade cue category (start, during, or end of the reach movement) on the durations of the directing and guiding phases. The duration of the directing phase (M = 511 ms, SE = 21 ms) did not depend on feedback (*F*(1,24) = 1.09; *p* = .31), cue category (*F*(2,48) = .92; *p* = .40), or the interaction (*F*(2,48) = .37; *p* = .69). The duration of the guiding phase was greater (*F*(1,24) = 7.58; *p* = .012) in the visual condition (M = 175 ms, SE = 20 ms) than in the visual-haptic condition (M = 111 ms; SE = 11 ms), but did not depend on cue category (*F*(2,48) = 1.01; *p* = .37) or the interaction between feedback and cue position (*F*(2,48) = 1.65; *p* = .20). These results indicate that participants did not alter the way they reached depending on when the saccade target was cued. In addition, whereas the final guiding, or approach, phase of the reaching movement depended on contact feedback, the initial larger amplitude directing phase was less affected.

Figure 4A shows, for each saccade cue condition, cumulative distributions of saccade onset times, relative to the end of the movement, for the visual and visual-haptic conditions. As expected, when the saccade cue was presented at the end of the movement, there was distribution of saccade onset times were similar for the two conditions. However, when the cue was presented at the start of the movement, saccade onset times were earlier for the visual-haptic condition compared to the visual condition. Note that the offset between these distributions is similar to the mean duration of the of the guiding phase in the visual-haptic condition (light blue regions). When the saccade cue was presented during the reach, saccade onsets were earlier for the visual-haptic condition for saccade initiated before the end of the hand movement, but saccade onsets were similar for the two conditions for saccades initiated after the end of the hand movement.

**Figure 4.**
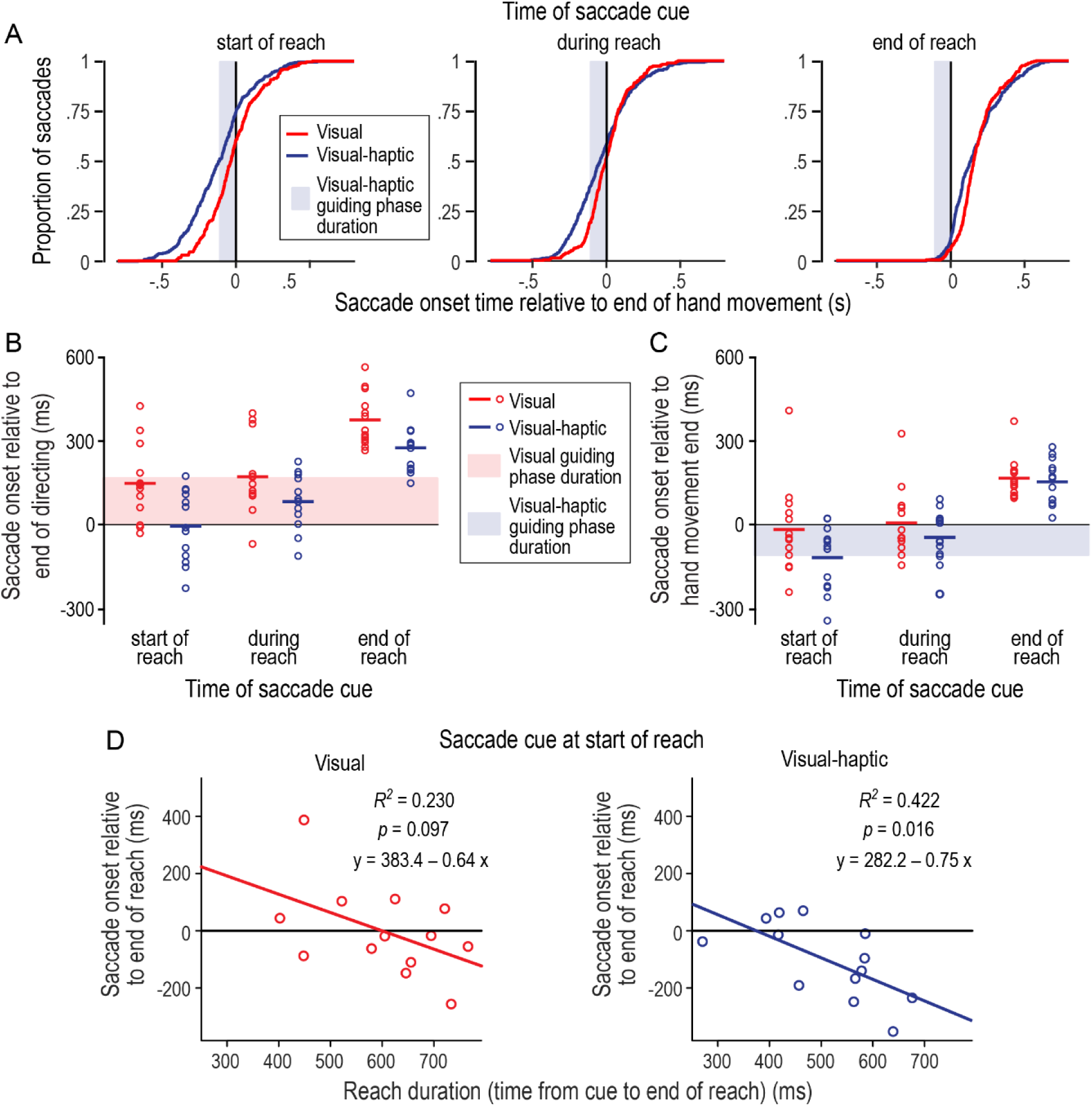
(A) Cumulative frequencies, for each saccade cue time, of saccade onset times, relative to the end of the hand movement, for the visual and visual-haptic conditions. The light blue regions represent the group mean duration of the guiding phase in the visual-haptic condition. The distributions include all trials from all participants. (B, C) Horizontal lines show, for each saccade cue category, group mean saccadic onset times relative to the end of the directing phase (B) and end of the hand movement (C) in the visual and visual-haptic conditions. Dots represent individual participant medians. The heights of the light red and blue regions represent the group mean durations of the guiding phase in the visual and visual-haptic conditions, respectively. (D) Relationship, across participants, between saccade onset relative to reach end and reach duration (time from cue at reach start to reach end) for the visual and visual-haptic conditions. Data for trials with the saccade cue at the start of reach.

Figures 4B and C show mean saccade onset—relative to the end of the directing phase and the movement, respectively—as a function of feedback condition and cue category. The effects of gaze anchoring are most clearly revealed when the saccade cue is delivered at the start of the reach. On average, in the visual condition saccade onset occurs at around the time of the end of the hand movement (M = −3 ms, SE = 44 ms), and in the visual-haptic condition saccade onset occurs at around the time of the end of the directing phase (M = 5 ms, SE = 37 ms). When the saccade cue is delivered at the end of the reach, the saccade onset occurred approximately 150 ms after the end of the hand movement and was similar for the visual (M = 146, SE = 15 ms) and visual-haptic (M = 137, SE = 15 ms) condition. When the saccade cue was delivered during the reach, the saccade onset in the visual condition was similar to when the cue was delivered at the start of the reach; on average, it occurred around the time of the end of the hand movement (M = 10 ms, SE = 36 ms). In contrast, in the visual-haptic condition, the saccade onset occurred before the time of end of the hand movement (M = −20 ms, SE = 29 ms), and approximate mid-way through the guiding phase. The observation that gaze was on average still anchored during the beginning of the guiding phase presumably reflects the limited time between the cue and the end of the direction phase in which to respond to the cue.

To quantify these observations, we assessed the effect of feedback condition and saccade cue category on gaze anchoring using a mixed-factor ANOVA. Consistent with the results shown in Figures F and G, the time of saccade onset relative to the time of reach end was significantly affected by feedback condition (*F*_1,24_ = 5.05; *p* = 0.03; *_ηη_* = 0.17), cue category (*F*_2,48_ = 637.85; p > .001; *_ηη_* = 0.96), and there was a significant interaction (*F*_2,48_ = 3.84; p = 0.03; *_ηη_* = 0.14).

Overall, these results provide evidence that gaze anchoring is not only observed in the visual condition, as expected, but also in the visual-haptic. However, gaze appears to be anchored to different events in these conditions. This is most clearly seen when the saccade cue is delivered at the start of the movement. Specifically, on average gaze appears to be anchored to the end of the movement in the visual condition, and to the end of the directing phase in the visual-haptic condition.

Despite the clear difference in gaze anchoring between conditions, we observed considerable variability in saccade timing across participants. This variability in gaze anchoring may arise, in part, because different participants tend to anchor on different time points—e.g., before or after the end of the directing phase, as we have defined it, in the visual-haptic condition. Moreover, our method of determining the change from the directing to the guiding phase is based on noisy movement data and thus a mere estimation.

Given the considerable variability across participants in saccade latency, we asked whether this variability is linked to variability in the reach movement. Focusing on trials in which the saccade cue was delivered at the start of the reach, we looked at the relationship between saccade onset relative to the end of movement and reach duration in the visual and visual-haptic conditions (Fig. 4D). In the visual-haptic condition, saccade onset, relative to the end of the reach movement, decreased with reach duration (*p* < 0.05); however, this trend was not significant in the visual condition. Given that the duration of the guiding phase is positively correlated with the overall reach duration (*r* = 0.67; *p* = 0.01), this result is consistent with the observation that saccades are, on average, anchored to the end of the directing phase and the end of the movement in the visual-haptic and visual conditions, respectively.

### Anchoring of gaze but not the hand in two-handed reaching

In the two-handed reaching task, we asked how the timing of the gaze shift to the secondary target would be affected if participants were asked to also move their left hand to the secondary target. More specifically, we asked whether gaze anchoring would still be observed and, if so, whether the left hand would also exhibit anchoring. Participants used their right hand to reach to the fixated right-hand reach target, and received only visual feedback about target contact (just as in the visual condition in the one-handed reaching task). The secondary—combined saccade and left hand—target was cued when the right hand reached 5, 50, and 98% of the distance to the target, instructing participants to move both their gaze (from the fixated reach target) and their left hand (from the left-hand start position) to this secondary target as soon as possible.

Figure 5A shows gaze, right hand, and left-hand paths and target positions for a representative trial in a screen-centred reference frame. Figure 5B shows gaze and hand velocities as a function of time for the same. The time of the cue and the end times of the directing phase and movement are shown by vertical lines. In this trial, the top potential saccade target was cued as the saccade target just after the right-hand movement started. Note that the left hand started moving towards the secondary target well before the right-hand movement ended, whereas gaze shifted to the secondary target at the end of the right-hand movement.

**Figure 5.**
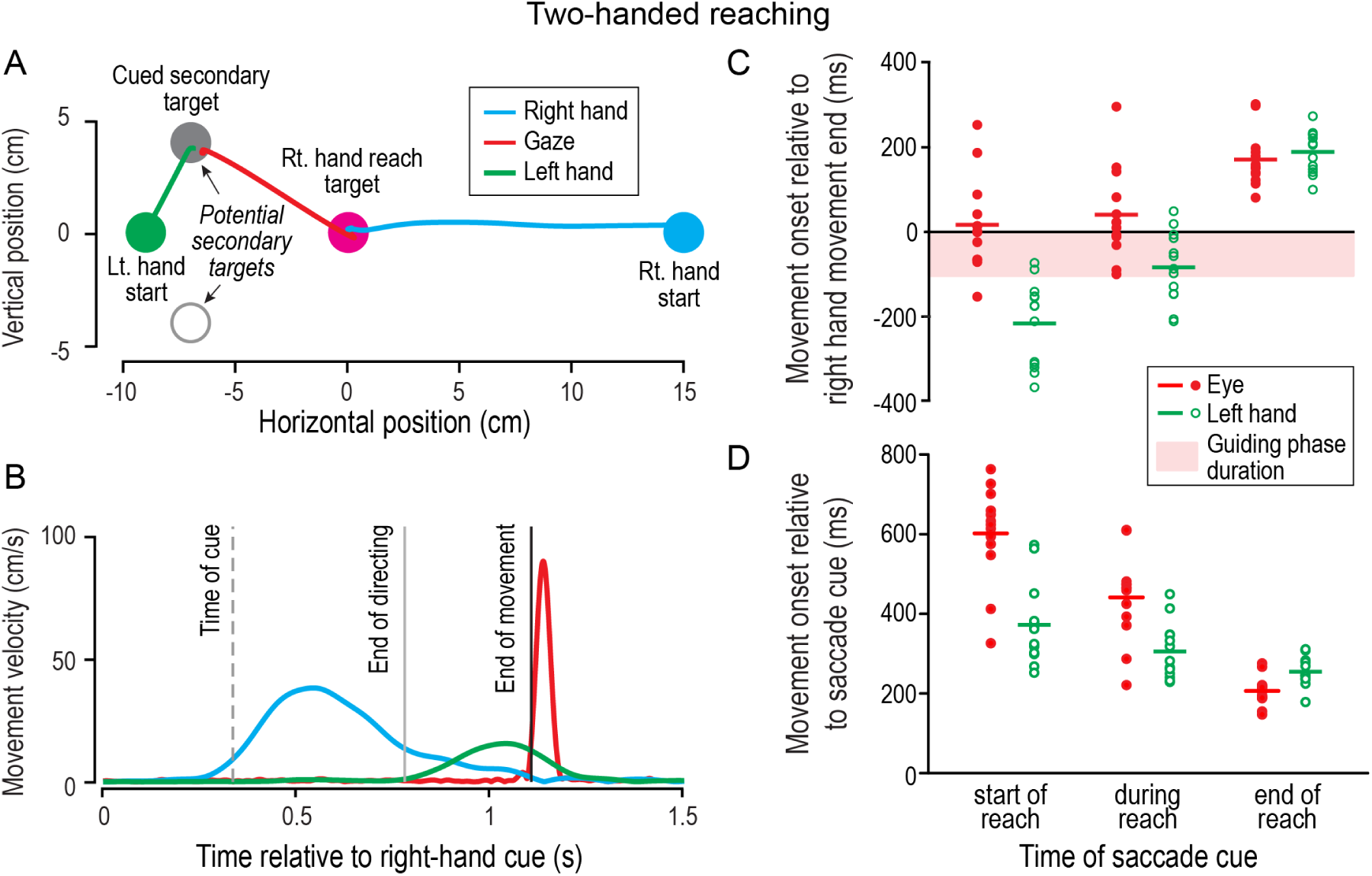
Results from the two-handed visually guided reaching task. (A) Gaze, left- and right-hand (i.e., cursor) paths, and target positions, in screen coordinates, from a representative trial. Participants used their right hand to reach to the right-hand reach target and were instructed to shift their gaze and move their left hand to the secondary target—randomly selected for two potential secondary targets—when it was cued during the right-hand movement. (B) Gaze and left and right-hand tangential velocity profiles from the trial in A. The times of the cue and ends of the directing phase and movement are indicated by vertical lines. (C, D) Horizontal lines show, for each saccade cue category, group mean saccadic and left-hand onset times relative to the end of the right-hand movement (C) or the time of the saccade cue (D). Dots represent individual participant medians. The height of the light red region represents the group mean duration of the guiding phase.

Figure 5C shows mean saccade and left-hand movement onset times, relative to the end of the right-hand movement, for each cue category. In line with the results of the one-hand reaching task, on average, saccade onset occurred close to the end of the reach when the saccade cue was delivered at the start of the reach (M = 17 ms, SE = 30 ms) or during the reach (M = 41 ms, SE = 30 ms), and well after the end of the right-hand reach when the saccade cue was delivered at the end of the reach (M = 171 ms, SE = 18 ms). In contrast, the onset of the left-hand movement o occurred well before the end of the right-hand reach when the cue was delivered at the start of the reach (M = −216 ms, SE = 28 ms) or during the reach (M = −83 ms, SE = 23 ms). Similar to saccade onset, the left-hand onset occurred well after the end of the right-hand reach when the cue was delivered at the end of the right-hand reach (M = 190 ms, SE = 14 ms). Thus, the onset of the left hand was not anchored to the end of the reach.

To quantify these differences, we tested the effects of effector (eye vs. hand) and saccade cue category on saccade latency relative to end of the right-hand reach using repeated measures ANOVA. The saccade latency relative to the end of the reach was affected by both effector (*F_2,24_*= 18.71; *p* < 0.001; *_ηη_* = 0.61) and cue condition (*F_2,24_*= 206.64; *p* < 0.001; *_ηη_* = 0.95). In addition, consistent with the different pattern of eye and left-hand movement latencies, there was a significant interaction (*F_2,24_*= 60.48; *p* < 0.001; *_ηη_* = 0.83).

The interaction between eye and left-hand latencies can be clearly seen in Figure 5D, which shows saccade and left-hand movement onsets relative to the time of the saccade cue. Separate one-way repeated measures ANOVAs indicated that both eye (*F_2,24_* = 128.8; *p* < 0.001; *_ηη_* = 0.91) and left hand (*F_2,24_* = 16.95; *p* < 0.001; *_ηη_* = 0.59) latency relative to the saccade cue were affected by saccade cue category. However, whereas saccade latency markedly increased as the remaining right hand reach distance (98, 50 and 5 % for cues delivered at the start, during, or end of the reach) increased, the left-hand latency only increased slightly as the remaining reach distance increased. This increase in left hand latency may reflect changes in sensorimotor processing during the right-hand reaching movement. Overall, these results indicate that gaze anchoring does not extend to hand anchoring.

### Comparison of saccadic latencies in the one- and two-handed reaching task

We observed considerable variability, across participants, in terms of saccade latency relative to the saccade cue. To test whether the latency magnitude was participant specific, for each saccade cue category we compared saccade latencies of the one- and two-handed reaching tasks. Figure 6 shows, for each saccade cue category, saccade latency in the two-handed task as a function of saccadic latency in the one-handed task. Each circle represents a participant and is based on the median latency across trials. Note that for these plots, we only used one-handed reaching trials in which the location of the secondary gaze target was in one of the two locations used in the two-hand reaching trials. In other words, we matched saccade directions. For all three cue categories, linear regression revealed a significant slope (p < 0.001) close to 1 with a high R^2^ value (≥ 0.919). These results indicate that participants exhibit highly consistent saccadic reaction times across tasks.

**Figure 6.**
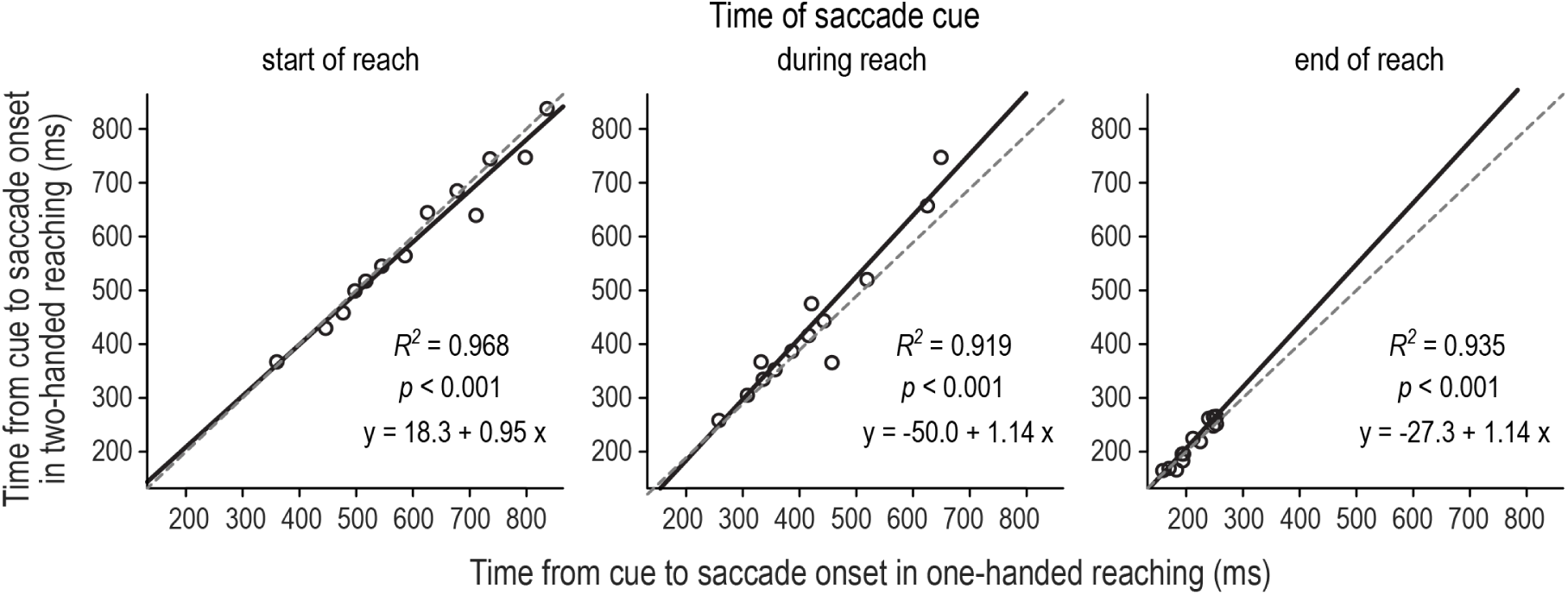
Relationship, across participants, between saccadic reaction times in the visual condition of the one-handed reaching task and the two-handed reaching task, with separate panels shown for each saccade cue category. Solid lines show best fit linear regression lines and the dash lines are unity lines.

## Discussion

The aim of this study was to investigate how quickly participants could initiate a visually-cued eye movement while engaging in manual reaching. In line with previous research, we found that saccades were delayed—relative to the cue—if the saccade was cued at the start or during the reach. However, the saccade delay depended on the availability of haptic feedback, with eye movements, on average, being initiated at the end of the reaching movement when feedback was only visual and at the end of the directing phase when participants received visual-haptic feedback upon contacting the reach target. We further tested the delay in movement initiation in a bimanual reaching task, in which the cued saccade target was also a left-hand reach target. Here, we found a similar saccade delay compared to one-handed reaching. However, the delay of the left-hand movement was much shorter than the saccade delay, indicating that the initiation of the left-hand reach was largely decoupled from gaze anchoring. Finally, we found that, in the visual condition, individual differences in how long saccades were delayed relative to the cue, were consistent across the one- and two-hand reaching tasks, suggesting that sensorimotor processing was similar in these two tasks.

### Gaze anchoring depends on available sensory feedback

In natural action tasks, humans coordinate their eye and hand movements in stereotypical ways (de Brouwer et al. 2021; Land 2006). When reaching for and manipulating objects, gaze commonly fixates action-relevant objects before the hand arrives, and then shifts to the next location of interest around the time that the hand contacts the target object (Ballard et al. 1992; Bowman et al. 2009; Johansson et al. 2001). Fixating the action goal has been shown to increase endpoint accuracy both when reaching toward and placing objects (Bock 1986; Desmurget and Grafton 2000; Fisk and Goodale 1985; Luabeya et al. 2024). Moreover, maintaining fixation at the reach goal appears to be linked to hand movement control. When a saccade target is flashed during a reaching movement, saccades to the flashed target are delayed compared to saccades that are cued after reach completion (Neggers and Bekkering 2000). The duration of this delay depends on the time at which the secondary target is flashed relative to reach completion, with saccades being delayed longer if the secondary target is flashed early during the reach (Neggers and Bekkering 2001). These results suggest that gaze anchoring occurs, at least in part, to allow central vision to be used towards the end of the hand movement to guide the hand to the target. What is not clear is whether gaze anchoring also occurs to allow peripheral vision and gaze-related signals (e.g., gaze proprioception) to be used earlier during the hand movement to direct the hand towards the vicinity of the target.

Here, we extend previous findings by showing that gaze anchoring not only occurs in the visual condition, in which central vision is required to guide the hand to the target via slow visual feedback loops, but also in the haptic condition, in which central vision is not required for this purpose. These results indicate that gaze anchoring can arise from visuomotor demands during the directing phase when reaching to the target. During the directing phase, peripheral vision of the hand can be used continuously to monitor the position of the hand and to rapidly correct for reach errors (Brenner and Smeets 1997; de Brouwer et al. 2018; Paillard 1996; Sarlegna et al. 2003; Saunders and Knill 2003, 2004). How much visuomotor control is needed when reaching towards a foveated target depends on task demands, such as the precision requirements at the reach goal (Illamperuma and Fooken 2024; Rand and Stelmach 2010; Sims et al. 2011), the time available to complete sequential reaches (Deconinck et al. 2011), the reward structure of the task (Abekawa et al. 2021), and whether people act in real or virtual environments (Lavoie et al., 2024; 2025). Thus, the strength of gaze anchoring is generally modulated by the structure of the action task and the environment. Yet, it should be noted that gaze anchoring—unlike more reflexive responses that tend to be tightly tied to a specific event—can vary considerably across participants.

Previous work has described the role of visual feedback in visually-guided reaching and grasping (Janssen and Scherberger 2015; Sabes 2000). However, in real-world action tasks, humans not only rely on visual but also multisensory signals to guide reach-to-grasp movements (Betti et al. 2021). For example, when grasping an object, tactile information from the fingertips is used to rapidly adjust hand kinematics and the force exerted to grasp the object (Johansson and Flanagan 2009; Pruszynski et al. 2016, 2018). Moreover, whether and when an object is foveated prior to grasping depends on the availability of haptic feedback upon contacting the object. Whereas manipulating objects with fingertips can be guided by tactile feedback, observers prefer to foveate objects until they are grasped when performing the manipulation task with a tool (Fooken et al. 2024b). Similarly, when people are deprived of haptic feedback—which is common in virtual reality—they spend more time looking at their current action and less time looking ahead to the next action site (Lavoie et al., 2025). Taken together, results from the current study and the literature highlight that saccade inhibition during reaching is modulated by the sensory information available to control the reaching movement.

### Dissociation between eye and hand movements in bimanual reaching

As highlighted in the previous section, the synergetic link between eye and hand movement control can be modulated by visuomotor task demands (de Brouwer and Spering 2022; Coudiere and Danion 2024; Epelboim et al., 1995; Sailer et al. 2000). For example, rewarding fast-latency saccades to a saccade target that is cued during a reaching movement increases the occurrence of non-anchored saccades (Abekawa et al. 2021). Here, we tested whether making the saccade target an additional, left-handed movement target would affect gaze anchoring in any way. We found that in the two-handed reaching tasks, saccades were delayed as long as in the one-handed reaching task. However, the initiation of left-handed reaches was to some degree decoupled from gaze anchoring. These findings are in line with research showing that the latencies of eye-and hand movements are only weakly correlated when humans reactively point to visual targets or rapidly respond to sudden changes in target position (Fooken et al. 2024a; Prablanc et al. 1979). Thus, our results provide further evidence that whereas sensory information is shared between the eye and hand movement system, movement initiation and execution may be controlled in parallel.

Neurophysiological studies have shown that the posterior parietal cortex plays a crucial role in coordinating eye and hand movements in visually-guided reaching (Andersen et al. 1997; Battaglia-Mayer et al. 2015; Buneo and Andersen 2006; Dean et al. 2012; Passarelli et al. 2021; Snyder et al. 2000). During reaching movements, neural firing in the lateral intraparietal area (LIP), an area associated with saccade and attentional control (Andersen et al. 1992; Barash et al. 1991; Bisley and Mirpour 2019), is systematically modulated by neural activity in the parietal reach region (PRR; Hagan et al. 2012; Hagan and Pesaran 2022). Specifically, saccades are transiently suppressed through a ‘reach-to-saccade communication channel’. One possibility is that whereas the reach target is represented in a common (oculomotor) reference frame (Batista et al. 1999; Carey 2000; Vesia and Crawford 2012), PRR and LIP operate in parallel to plan and control movement execution (Kang et al. 2024). Overall communication between PRR and LIP may be functionally organized such that the suppression of saccades during reaching supports goal-directed behaviour.

### Individual differences in saccade latency

The tendency to inhibit saccades shortly before action-relevant events has not only been described in laboratory studies but also in the wild. Expert performance in many targeted action tasks, such as golf putting or basketball free throw, is characterized by a systematic suppression of saccades before the action is executed, a phenomenon known as ‘quiet eye’ (Vickers 1992, 1996). Maintaining steady fixation of the action goal is thought to facilitate information and attentional processing and aid motor preparation (Gonzalez et al. 2017; Vickers 2007). Yet, more dynamic action tasks require a disengagement of the fixation to make gaze available to gather new visual information and support ongoing action control. We observed that how long saccades were delayed during the ongoing reach greatly varied between individuals, reiterating the observation that eye movement behaviour systematically differs between individuals (Bargary et al. 2017; Castelhano and Henderson 2008; de Haas et al. 2019). Interindividual eye movement differences even persist across different tasks, if similar sensorimotor demands are required (Goettker and Gegenfurtner 2024). We observed that the delay in saccade latency to the visually cued target was consistent within individuals irrespective of whether participants performed the one- or two-handed reaching task. Taken together, these results suggest that the tradeoff between perceptual and motor processing is a balance that is tailored to each individual’s sensorimotor ability. Although these sensorimotor traits may to some degree be hardwired (Kennedy et al. 2017), the fact that eye movement patterns in visual and motor experts are similar between individuals suggest that visuomotor experience also plays a role (Reingold and Sheridan 2011; Vickers 2007).

## Conclusion

The current study investigates mechanisms of gaze anchoring, a phenomenon that describes the inability to move the eyes away from the reach goal to a visually cued target while the reaching movement is ongoing. We found that how long saccades were delayed—relative to the time of saccade cue—depended on the type of feedback participants received upon contacting the reach target. Specifically, whereas gaze anchoring was linked to the end of the reaching movement when the target was only visual, gaze anchoring was linked to the end of the directing phase when haptic information was available upon contact between the hand and the reach target. We further found that gaze anchoring did not depend on whether the cued target was only a saccade target or a combined saccade and left-hand target, and gaze anchoring was not systematically related to the initiation of a simultaneous left-handed movement. Effects of gaze anchoring were highly consistent within individuals and correlated across tasks. Taken together, our results suggest that visual feedback is continuously used to support goal-directed action until other sensory feedback is available or the action goal is attained. However, while the eyes are anchored to the reach target, a secondary goal-directed movement can be planned and initiated in parallel. The timing of these interacting visuomotor control mechanisms appears to be individual-specific and may indicate differences in trading off perceptual and sensorimotor processes.

### Citation diversity statement

Recent work in several fields of science has identified a bias in citation practices such that papers from women and other minority scholars are under-cited relative to the number of such papers in the field (Bertolero et al. 2020; Caplar et al. 2017; Chatterjee and Werner 2021; Dion et al. 2018; Dworkin et al. 2020; Fulvio et al. 2021; Maliniak et al. 2013; Mitchell et al. 2013; Wang et al. 2021). Here we sought to proactively consider choosing references that reflect the diversity of the field in thought, form of contribution, gender, race, ethnicity, and other factors. First, we obtained the predicted gender of the first and last author of each reference by using databases that store the probability of a first name being carried by a woman (Dworkin et al. 2020; Zhou et al. 2020). By this measure (and excluding self-citations to the first and last authors of our current paper), our references contain 15.28% woman(first)/woman(last), 10.27% man/woman, 19.44% woman/man, and 55.01% man/man. This method is limited in that a) names, pronouns, and social media profiles used to construct the databases may not, in every case, be indicative of gender identity and b) it cannot account for intersex, non-binary, or transgender people. Second, we obtained the predicted racial/ethnic category of the first and last author of each reference by databases that store the probability of a first and last name being carried by an author of colour (Ambekar et al. 2009; Chintalapati et al. 2023). By this measure (and excluding self-citations), our references contain 4.62% author of colour (first)/author of colour(last), 17.34% white author/author of colour, 18.30% author of colour/white author, and 59.74% white author/white author. This method is limited in that a) names and Florida Voter Data to make the predictions may not be indicative of racial/ethnic identity, and b) it cannot account for Indigenous and mixed-race authors, or those who may face differential biases due to the ambiguous racialization or ethnicization of their names. We look forward to future work that could help us to better understand how to support equitable practices in science.

## Credit authorship contribution statement

Conceptualization (JF, JRF), Methodology (JF, JRF), Validation (JF, JRF), Formal analysis (JF, JRF), Data Curation (JF, NI), Writing: Original Draft (JF, NI), Writing: review & editing (JF, NI, JRF, JPG), Visualization (JF, JRF), Supervision (JF, JRF, JPG), Funding acquisition (JF, JRF, JPG)

## Acknowledgements

This work was supported by a Deutsche Forschungsgemeinschaft (DFG) Research Fellowship to JF (FO 1347/1-1), an NSERC Discovery grant and a Canadian Institutes of Health Research grant (RGPIN/05944-2019) to JRF, and operating grants from the Canadian Institutes of Health Research Grant (PJT175012) and the Natural Sciences and Engineering Research Council (RGPIN-2017-04684; RGPIN-2024-06773), awarded to JPG. The authors would like to thank Martin York for his help with coding the experiment routine. We also would like to thank Elissa Robichaud and Bethany Piekkola for their help with participant recruitment as well as data collection and preprocessing.

